# Genome-wide CRISPR knockout screen reveals the landscape of essential genes across the porcine genome

**DOI:** 10.1101/2025.02.28.640817

**Authors:** Thomas Harvey, Victor Boyartchuk, Maren van Son, Arne B. Gjuvsland, Eli Grindflek, Matthew Kent

## Abstract

Characterization of essential genes across the genome is fundamental to understanding cellular functions at a molecular level. While significant progress has been made in characterizing essential genes in human and mouse models, relatively little is known about essential genes in the porcine genome. Pigs are an important production species and are now emerging as valuable models for studying human diseases due to their physiological similarities to humans. To map essential genes across the porcine genome, we have developed a novel porcine genome-wide CRISPR knockout screening library (pGeCKO) and applied it to two porcine cell lines, PK15 and IPEC-J2. We identified 2,245 essential genes in PK15 cells and 919 essential genes in IPEC-J2 cells, with 683 of these shared between both cell lines. Functional analyses revealed that most essential genes are involved in core cellular processes such as cell cycle regulation, DNA replication, transcription, and translation. Comparative analysis with human essential genes from the DepMap project revealed that over half of the genes are shared with humans and the rest are porcine-specific. These porcine-specific essential genes included genes in core functional pathways related to protein and RNA processing as well as many related to N-glycan biosynthesis, signal transduction, and several long-noncoding RNAs. This work provides a new resource for leveraging porcine models in disease research, enhancing our understanding of porcine genetics and its implications for human health.

## 2 Introduction

Cellular fitness relies on the complex interaction of thousands of gene products. Studies using human cells (Tsherniak et al., 2017) have revealed a complex landscape of gene essentiality, with a core group of genes absolutely necessary for cellular function and a highly context dependent group of genes that are only necessary under certain conditions, developmental stages, tissues, or genetic backgrounds (Dvir et al., 2022). In pigs, little is known about gene essentiality, so much is assumed based on studies in other mammalian species. Given the context dependent nature of essential genes (Larrimore & Rancati, 2019) and around 90 million years of evolutionary divergence (Groenen et al., 2012), many of these assumptions may not hold. Therefore, it is critical to understand the landscape of gene essentiality within a pig specific model. Given the physiological similarities of pigs and humans, pigs are increasingly recognized as valuable models for studying human diseases (Hou et al., 2022). In recent years, pig models have advanced our understanding of several diseases with high societal impact, including myocardial infarction (Hobby et al., 2019; Yuan et al., 2020), renal disease (Rodríguez et al., 2020), non-alcoholic fatty liver disease (Wang et al., 2020), and congenital hypothyroidism (Zhang et al., 2017). Cancer biology is one field that could greatly benefit from porcine models as treatments developed in pigs may translate more effectively to the clinic than treatments developed in mouse models (Mak et al.; Saur & Schnieke, 2022). The recent development of the Oncopig cancer model is a promising step towards closing this species gap (Joshi et al., 2024). Pigs are also a promising source of healthy organs for transplantation, with xenotransplantation of humanized pig organs already attempted, albeit with limited success so far (Anand et al., 2023). To fully harness the potential of porcine models in studying and treating human diseases, we need a better understanding of porcine genetics, including their similarities and differences to humans. Identifying pig-specific essential genes will be crucial for elucidating species-specific molecular physiology and preventing the disruption of key fitness genes during genetic modification.

Extensive research has been conducted on essential gene characterization in human and mouse models (Bartha et al., 2018; Cacheiro et al., 2020; Liang et al., 2024; Wang et al., 2015). Traditionally, researchers have relied on genome-wide loss-of function screening studies using RNA interference (Kampmann et al., 2015). More recently, genome-wide CRISPR knockout (GeCKO) screens (Shalem et al., 2014) have emerged as a more effective approach for identifying essential genes (Evers et al., 2016) and have proven to be an extremely useful tool in modern cancer research (Behan et al., 2019). GeCKO screening is particularly well suited to identification of essential genes because it generates a population of cells, each with a single gene knocked out, collectively covering all genes in the genome. Over time, genes that are critical to cell function (essential genes) will naturally drop out of the cell population, providing an unbiased, genome-wide portrait of gene essentiality. One major initiative, Project Achilles (Tsherniak et al., 2017), aims to identify essential genes across all cancer types to uncover cancer specific genetic dependencies and develop targeted therapies for specific cancer types. To date, the project has performed over 1000 GeCKO screens across more than 30 cancer cell lineages. Similarly, the International Mouse Phenotyping Consortium have tested gene essentiality *in vivo* by producing knockout mice and comparing results with human cell line screens (Cacheiro et al., 2020). These studies have revealed that many genes that are non-essential for cell survival are essential for development. Moreover, they highlight key differences in essential gene requirements between humans and mice, emphasizing the need for species-specific investigations of gene essentiality.

For the past decade, GeCKO screening has been a critical technology for studying gene essentiality and host-pathogen interactions, however most studies in higher eukaryotes have been limited to human cell lines. Recently, screens have been successfully conducted in several farmed animals to identify host genes controlling susceptibility to industrially relevant pathogens, including four in pig (Jiang et al., 2022; Shen et al., 2024; Zhao et al., 2020; Zhou et al., 2021), one in cattle (Tan et al., 2023), and one in chicken (Jin et al., 2025). The expansion of this technology to key agricultural species opens new possibilities for selective breeding and genome modification to enhance disease resistance, improving sustainability and animal welfare in livestock production.

In this study, we develop a novel pig specific GeCKO screening library (pGeCKO) and use it to identify essential genes in two porcine cell lines. We then compare this list of genes to essential genes in human cells to determine shared and pig-specific essential genes. Additionally, we identify several essential long noncoding RNAs (lncRNAs) with unknown function. The goal of this work is to; (i) identify essential genetic elements in pig to support future research in disease modeling, xenotransplantation, and genetic modification, and (ii) validate our pig specific GeCKO screening library as a robust framework for future studies on pig diseases.

## 3 Materials and Methods

### 3.1 Cell culture

HEK293T cells were obtained from the Centre of Molecular Inflammation Research (CEMIR, NTNU, Norway) and maintained in DMEM (Merck) with 10% fetal bovine serum (Gibco), penicillin-streptomycin (ThermoFisher Scientific), and 2 mM L-glutamine (ThermoFisher Scientific). Intestinal porcine epithelial cells (IPEC-J2) were kindly provided by the Foods of Norway project (NMBU, Ås, Norway). Porcine kidney cells (PK15) cells were purchased from ATCC (CCL-33). Both porcine cell lines were maintained in DMEM (ThermoFisher Scientific) supplemented with 10% fetal bovine serum (ThermoFisher Scientific) and penicillin-streptomycin (ThermoFisher Scientific). All cells were cultured at 37 °C with 5% CO2 and subcultured at a ratio of 1:5 when they reached 70-80% confluency.

### 3.2 Lentiviral packaging and titration

Four T225 flasks were each seeded with 1.8×10^7^ HEK293T cells 24 hours before transfection. Each flask was co-transfected at a molar ratio of 1:1.5:2 with pMD2.G (addgene #12259): psPAX2 (addgene #12260): pGeCKO library. Specifically, lipofection was performed as follows: 15.3 μg pMD2.G, 23.4 μg psPAX2, and 30.6 μg pGeCKO library was added to 2.25 mL OptiMEM (ThermoFisher Scientific); 297 μl PLUS reagent (ThermoFisher Scientific) was added to 2.25 mL OptiMEM; and 270μL lipofectamine 2000 was added to 4.5 mL OptiMEM. The PLUS reagent was added to the DNA solution, inverted several times, and incubated at room temperature for 5 minutes. The DNA and PLUS mixture was then added to the lipofectamine solution, inverted several times, and incubated at room temperature for 5 minutes. The total of 9 mL of resulting lipid-DNA complexes was slowly added to each T225 flask containing 45 mL of culture media, gently mixed, and placed at 37 °C, 5% CO2. After four hours media was replaced with 45 mL fresh culture media and transfected cells were incubated at 37 °C, 5% CO2 for 3 days. Lentiviral particles were harvested in the combined supernatant (180 mL total) as follows; supernatant was passed through a 0.45 μm syringe filter (Merck) to remove cells and other large particles before being concentrated using a 100 kDa cutoff column (ThermoFisher Scientific). Aliquots of 15 mL supernatant were loaded onto six 100 kDa cutoff columns and centrifuged at 1500xg for 30 minutes in a swinging bucket rotor. Centrifugation was repeated using the same columns but applying a second volume of 15 mL supernatant. Virus retained by the column was then washed by resuspending the concentrated viral suspension in 15 mL PBS, centrifuging at 1500xg, and repeating for two washes in total. Concentrated virus was brought up to a final volume of 2 mL with PBS, pooled, split into 1 mL aliquots, and stored at –80 °C.

Functional lentivirus titer was determined empirically for each cell line by reverse transduction. Serial 2-fold dilutions of lentivirus were added in duplicate to an equal volume of cells at a concentration of 1.5×10^5^ cells/mL. Then, 2 mL of transduced cells was added to 2 wells of a 6-well plate for each virus dilution and placed at 37 °C, 5% CO2. After 24 hours, virus containing media in one of the wells was replaced with growth media, and the other was replaced with growth media containing 2 μg/mL puromycin. After 72 hours of selection, cells in both wells were trypsinized and counted. Multiplicity of infection (MOI) for each virus dilution was calculated as follows:

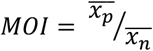

Where 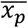 is the average cell count of puromycin selected cells, 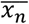 is the average cell count of non-selected cells. Viral titer in transduction units per mL (TU/mL) was calculated for the dilution with MOI closest to 0.3 as follows:

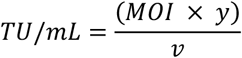

Where y is the number of cells seeded in the well, and v is the volume of virus in mL.

### 3.3 pGeCKO library design

All possible 20 nucleotide long sequences adjacent to an NGG Cas9 PAM sequence were identified within all coding and a subset of non-coding transcripts annotated in the porcine genome assembly 11.1. For each guide the number of additional binding sites in the porcine genome that are a perfect match (MM0) or have one or more mismatched nucleotides (MM1, MM2, etc) was calculated. Guide sequences were excluded if they contained homopolymers greater than 5 nucleotides. For each gene, all guides with MM0≥1 were discarded if there were at least 20 guides remaining. Cutting efficiency for guides targeting coding genes was calculated using the complete Doench scoring model that considers the location of the cut in the protein coding portion of the sequence (Doench et al., 2016). For each gene, individual guides were ranked based on the cutting efficiency, number of additional perfect and imperfect binding sites, and location of the cut within the coding sequence. The top 5 guides for each coding gene and lncRNA transcripts were included in the library. For a subset on non-coding RNAs, 3 guides were included in the library. Following assembly, the library was further filtered to remove redundant sequences. The final library contained 120,351 unique elements targeting 26,796 genomic features of which 19,746 were annotated protein coding genes. The library was synthesized by GENEWIZ and cloned into pLentiCRISPR v2 vector (Sanjana et al., 2014). Representation of guides in the final cloned library was confirmed by Illumina sequencing.

### 3.4 GeCKO screening

#### 3.4.1 IPEC-J2 cells

Genome-wide knockout cell populations were prepared by reverse transduction of IPEC-J2 cells with the pGeCKO lentiviral library. In short, 1.68×10^8^ IPEC-J2 cells were resuspended at a concentration of 4×10^5^ cells/mL in complete culture media (420 mL) containing 8 μg/mL polybrene and combined with an equal volume of lentivirus at a concentration of 1.2×10^5^ TU/mL in polybrene containing media (MOI of 0.3). Cells were mixed by inversion and seeded into 21 T175 flasks at a density of 4.6×10^4^ cells/cm^2^ then placed in a 37 °C, 5% CO2 incubator. An untransduced control sample of IPEC-J2 cells was also included by performing a mock transduction using polybrene containing media only. After 24 hours lentivirus containing media was replaced with 40 mL culture media containing 2 μg/mL puromycin and placed at 37 °C, 5% CO2. Media was changed every 3 days and cells were subcultured into a minimum of 2 flasks as needed every 3-4 days at a density of 1.4×10^4^ cells/cm^2^. Samples of at least 1×10^7^ cells were taken 3-, 10-, and 26-days post transduction.

#### 3.4.2 PK15 cells

Single gene knockout populations of PK15 cells were prepared as described above with the following exceptions. Reverse transduction was performed using 1.8×10^8^ cells at a concentration of 5×10^5^ cells/mL and lentivirus at a concentration of 1.5×10^5^ TU/mL (MOI of 0.3) to achieve a seeding density of 5.7×10^4^ cells/cm^2^. Cells were maintained by subculturing a minimum of 6 flasks as needed at a seeding density of 4.6×10^4^ cells/cm^2^. Samples were taken 5-, 13-, 21-, and 26-days post transduction.

### 3.5 Guide amplicon library preparation and sequencing

Genomic DNA was extracted from a maximum of 2×10^7^ cells using the Quick-DNA Midiprep Plus Kit (Zymo Research) according to manufacturer protocols. gDNA concentration was determined using the Qubit broad range kit (Invitrogen) and quality was assessed by gel electrophoresis and TapeStation genomic kit (Agilent). Amplicon libraries for next generation sequencing were prepared by PCR amplification from a maximum of 23 μg gDNA using P5 ARGON and P7 BEAKER Broad Institute hybrid primers composed of Illumina cell attachment and sequencing primer sites (Table S1). In addition, P5 primers contained a stagger region of different length and P7 primers contained Broad Institute barcodes for deconvolution after pooled sequencing (Table S1). The last 4 bases of the 3’ end of all primers were phosphorothioated to prevent exonuclease activity of proofreading polymerases. PCR amplification of each sample was split into 50μl reactions containing up to 1.5 μg gDNA, 1x NEBNext Ultra II Q5 master mix (NEB) and 0.5 μM of each primer. PCR reactions were run at 98 °C for 3 minutes, 25 cycles of 98 °C for 10 seconds, 56 °C for 10 seconds, 72 °C for 25 seconds, followed by a final extension of 72 °C for 2 minutes. PCR reactions of the same sample were pooled and cleaned up by 2 rounds of AMPure XP bead (Beckman Coulter) purification according to manufacturer protocols with a 1:1 ratio of beads to sample and 20 μl elution volume. Samples were sequenced on the NovaSeq 6000 (Illumina) with a minimum depth of 240 million reads per sample (PE150).

### 3.6 Data analysis

To assess the representation and relative abundance of DNA encoding each of the 118k gRNAs raw fastq reads were mapped and counted directly using MAGeCK (Li et al., 2014). Counts were normalized using the 1000 non-targeting control gRNAs. Essential genes were identified using the maximum likelihood estimation (MLE) module of MAGeCK with sampling time in weeks as a factor.

Essential gene results from pig cell lines were compared to human essential genes identified through the DepMap project (https://depmap.org/portal). Human essential genes were identified through CRISPR based screens across 56 kidney cell lines, 43 intestine cell lines, and all 1959 human cell lines available. Genes were considered essential if they had a dependency probability score above 0.5 in more than half of the cell lines screened.

## 4 Results

### 4.1 Essential genes across the porcine genome

To identify essential genetic elements in the porcine genome, we conducted genome-wide CRISPR screens using two porcine cell lines, PK15 and IPEC-J2, and looked for changes in gRNA abundance over a 26-day period (Figure 1a). Our porcine genome-wide knockout (pGeCKO) library comprised 120,351 guide RNAs (gRNA) targeting 19,746 coding genes (five guides per gene), 6,673 long-noncoding RNA (three guides per gene), and 337 miRNA (3 guides per gene). Additionally, it included 1000 non-targeting control gRNAs (Figure 1b), designed to have no effect on cell fitness. To assess gRNA dropout during library production and confirm actual composition of the pGeCKO library we performed amplicon sequencing of the plasmid library. Sequencing of gRNA-containing amplicons recovered from transduced cell populations at various time points revealed a gradual divergence from the initial gRNA distribution over time in both cell lines (Figure 1c).

**Figure 1:**
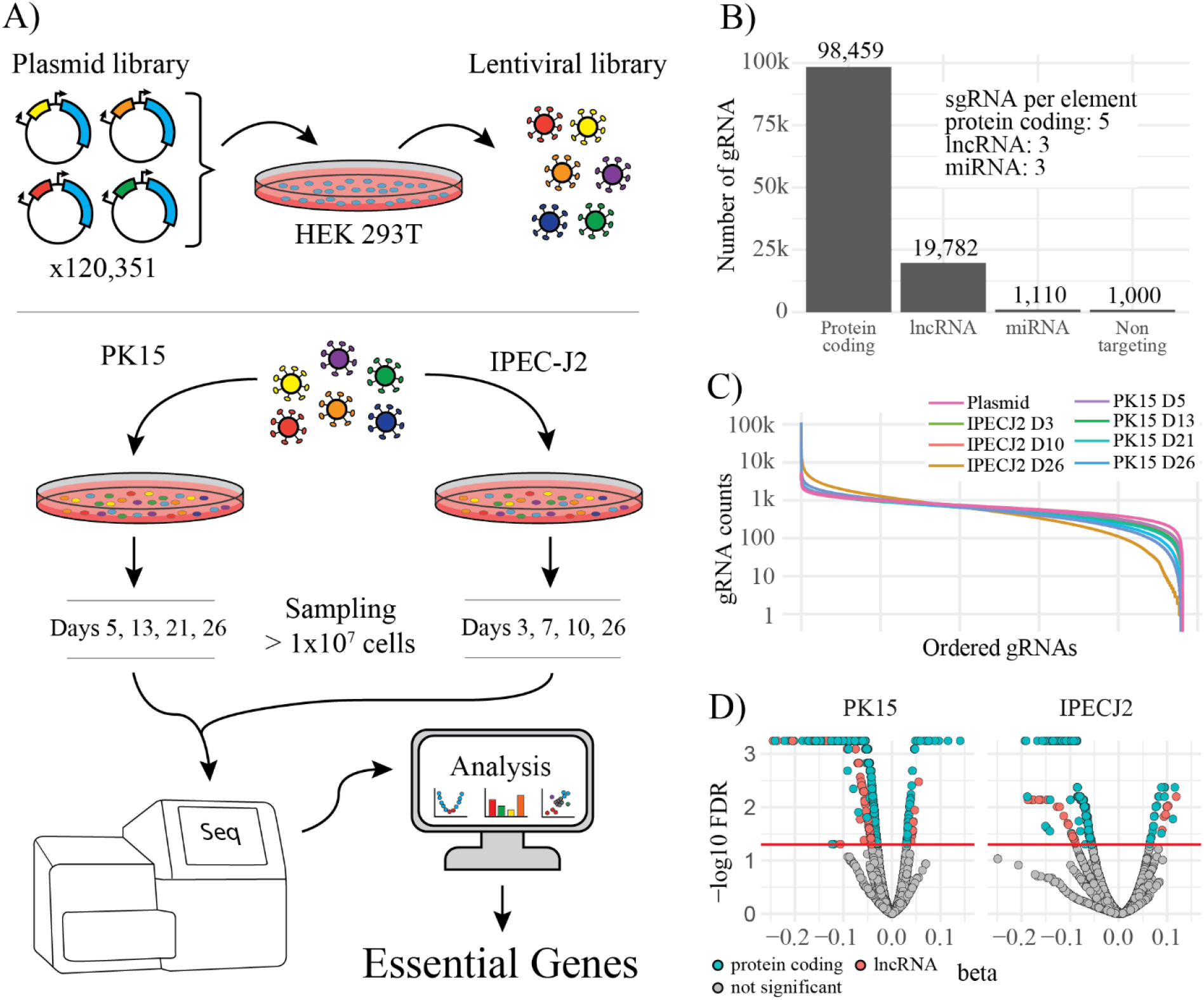
GeCKO library composition and performance. A) Flow chart of the GeCKO experiment. B) Composition of guide RNA targets in the pGeCKO library. C) Distribution of recovered guide counts in surviving cell populations at each sampling point. D) Identification of over– or under-represented gene targets in each cell line. Color corresponds to protein coding genes (green) or lncRNA (red).

Based on a model considering guide abundance across all four timepoints, we identified gRNAs targeting a total of 2,245 (PK15) and 919 (IPEC-J2) genes that clearly changed in abundance over time (FDR < 0.05) in PK15 and IPEC-J2 cells, respectively (Figure 1d, Table 1). Although there was evidence of both overrepresentation (indicating accelerated cell division) and underrepresentation (indicating decelerated cell division or cell death) of gRNAs, most significant changes were gRNAs that progressively decreased over the timeframe of the experiment. The majority of depleted guides were targeting either protein coding genes (2,048 PK15, 855 IPEC-J2) with the remainder targeting lncRNAs (118 PK15 and 26 IPEC-J2). Together genes in this set constitute putative essential genes since the absence of gRNA in the cell population indicates a loss of cells harboring these edits. We also identified a smaller subset of genes with corresponding gRNAs that increased in abundance over time (protein coding genes – 68 in PK15 and in 29 IPEC-J2; lncRNA genes – 11 in PK15 and 9 in IPEC-J2).

**Table 1:** Number of genes corresponding to decreasing or increasing gRNA abundance over time and number shared.

| Cell line | Down |  | Up |  | Total |
| --- | --- | --- | --- | --- | --- |
|  | Gene | lncRNA | Gene | lncRNA |  |
| PK15 | 2,048 | 118 | 68 | 11 | 2,245 |
| IPEC-J2 | 855 | 26 | 29 | 9 | 919 |
| Shared | 664 | 11 | 5 | 0 | 683 |

### 4.2 Functional characteristics of essential genes

To characterize the functions of genes identified in our study, we performed Kyoto encyclopedia of genes and genomes (KEGG) enrichment analysis on the enriched and depleted gene sets from both cell lines. Among the 2,481 essential genes identified, 683 genes were shared between the two cell lines, while 236 were unique to IPECJ2 and 1,562 were unique to PK15 (Figure 2a). KEGG analysis of each of these groups revealed that essential genes were generally enriched in five main pathway categories: ‘Cell cycle’, ‘Folding, sorting and degradation’, ‘Replication and repair’, ‘Transcription’, and ‘Translation’ (Figure 2a).

**Figure 2:**
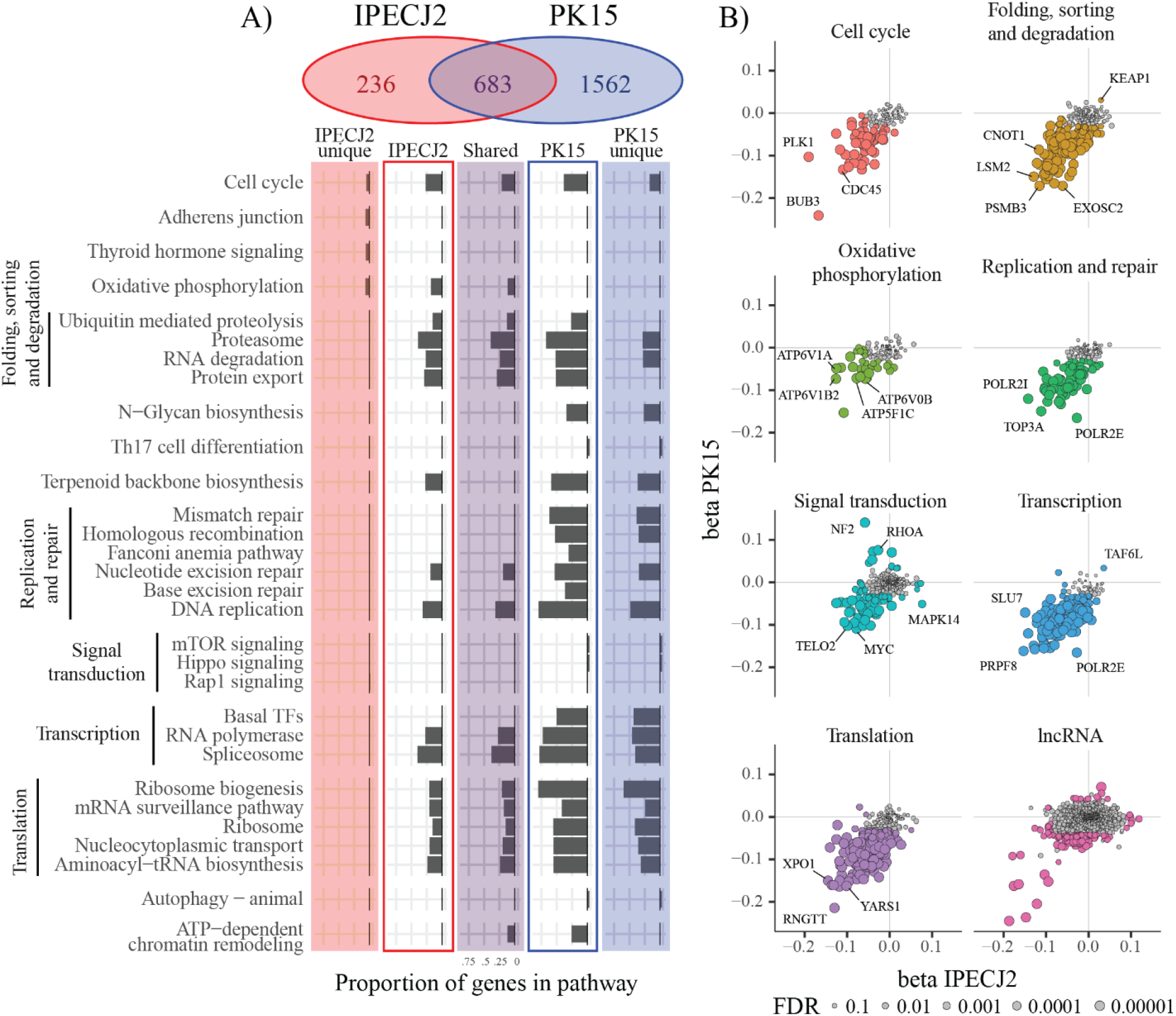
Pathway enrichment of pig essential genes. A) KEGG enrichment of essential genes in IPECJ2 cells (red border), PK15 cells (blue border), unique to each cell line (red and blue background), and shared between cell lines (purple background). B) Genes in several enriched pathway groups plotted by beta scores. Colored points are statistically significant (FDR < 0.05) in at least one cell line. Size corresponds to FDR of the most significant across the two cell lines.

Most genes identified in these pathways are well known to be involved in core cellular processes. For example, essential genes involved in cell cycle regulation included several key genes in mitotic spindle formation, such as BUB3 mitotic checkpoint protein (*BUB3*) and polo-like kinase 1 (*PLK1*). Additionally, five of the six genes of the origin recognition complex (*ORC1, ORC3, ORC4, ORC5*, and *ORC6*) were classified as essential in PK15 cells. The ‘Folding, sorting and degradation’ pathway contained 287 genes involved in ubiquitin mediated proteolysis, proteasome, RNA degradation, and protein export of which 64 were essential in both cell lines. Highly essential genes in this group included 17 components of the 26S proteasome complex and all 11 components of the nuclear exosome complex *(EXOSC1-10, DIS3*). We also identified many key members of the DNA replication complex, including genes of the DNA polymerase α-primase (*POLA1, PRIM2*), polymerase δ complex (*POLD1, POLD2, POLD3*), polymerase ε complex (*POLE*), and all members of the helicase complex (*MCM2-7*). Key components of transcription included seven RNA polymerase I, nine RNA polymerase II, and eight RNA polymerase III genes. Highly significant essential genes in translation pathways included the genes for mRNA capping (*RNGTT*), nuclear export (*XPO1*), and t-RNA synthetase (*YARS1*).

Guides that were enriched in GeCKO cell population (i.e. resulting in faster growth) targeted genes that were mostly not annotated to specific pathways (48 genes, Figure S1). Of the pathway annotated genes, most belonged to signal transduction pathways (14 genes, Figure 2b). Notably, this included known tumor suppressor genes *NF2* (Xu et al., 2024), *RHOA* (Dopeso et al., 2018), *TSC1* and *TSC2* (Kang et al., 2011).

Throughout our analysis we identified long noncoding RNAs (lncRNAs), a class of molecule gaining attention for their regulatory roles, in both enriched and depleted GeCKO populations. While most lncRNAs identified in our screen are cell line specific, the most significant enriched and depleted lncRNAs were common to both cell lines (11 genes, Figure 2b). The accession numbers of these essential lncRNAs are listed in Table S2.

### 4.3 Comparison of essential genes between pig and human

In order to validate our list of essential genes, and to find those that are unique to pig, we compared our pig essential gene list to those identified in human through the DepMap project (Tsherniak et al., 2017). Of 2,124 genes with a pig-human ortholog that were essential in either species, 1,149 genes (54.1%) were essential in both, 644 were essential in pig only (30.3%), and 331 were essential in human only (15.6%) (Figure 3a). Among the 575 shared essential genes involved in any pathway, the majority were associated with core cellular processes, including ‘Folding, sorting, and degradation’ (23.8%), ‘Transcription’ (17.5%), and ‘Translation’ (35.1%). Essential genes specific to pig included 455 in PK15 cells, 120 in IPECJ2 cells, and 69 in both cell lines. Notable genes in this category include 12 ribosomal genes, nine genes involved in protein processing in the endoplasmic reticulum, nine genes in RNA degradation, eight genes in aminoacyl-tRNA biosynthesis, eight genes involved in N-Glycan biosynthesis, and 70 genes involved in signal transduction.

**Figure 3:**
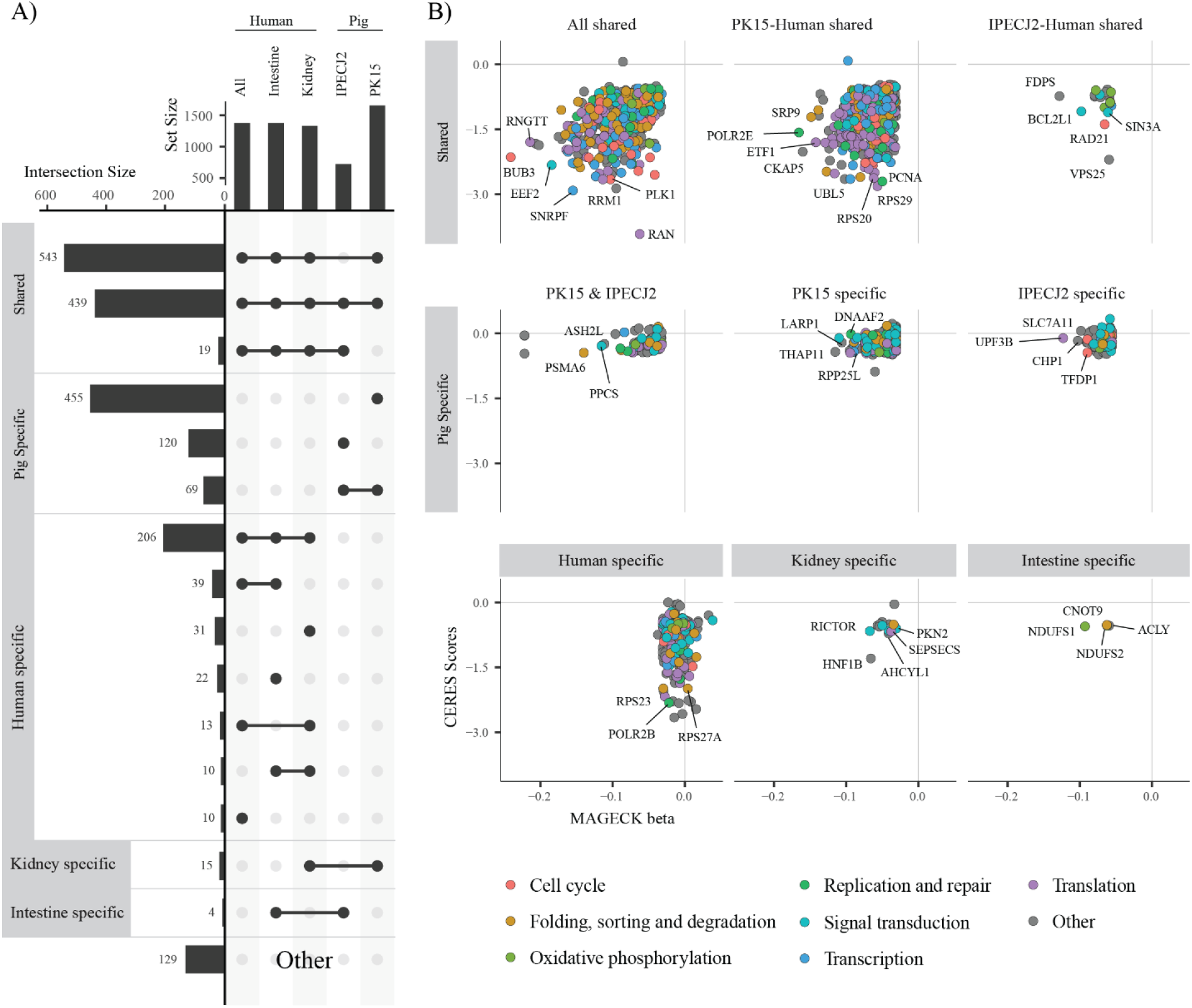
Comparison of essential genes between pig and human. A) Intersection of essential genes across screens in PK15, IPECJ2, human kidney, human kidney, and all human cell lines available in the DepMap project. B) Pig and human essential genes across intersection groups plotted by enrichment scores. Pig genes (x-axis) are plotted by MAGeCK beta scores and human genes (y-axis) are plotted by CERES scores. Scores from the PK15 and all human screens are used except for intersection groups for IPECJ2 where scores from the IPECJ2 screen are used and intestine specific where IPECJ2 and human intestine scores are used. Color corresponds to pathway membership of the gene.

Our analysis also identified a set of tissue specific essential genes that are conserved across species (i.e. essential in kidney or intestine in both human and pig). This set included 16 essential genes that were shared across species in kidney only. Two notable genes in this category include *HNF1B*, which is involved in nephron development (Massa et al., 2013; Sánchez-Cazorla et al., 2024), and *RICTOR*, which regulates cell growth and survival through the mTOR pathway (Jebali & Dumaz, 2018). In intestine cells, only four genes were essential across both species. Notably, two of these are members of the complex I mitochondrial respiratory chain 75-kD subunit, *NDUFS1* and *NDUFS2*. These genes are involved in NADH oxidation in the mitochondria, and mutations in these are found to be associated with complex I deficiency in humans (Bénit et al., 2001; Loeffen et al., 2001).

### 4.4 Varying rates of gRNA depletion

To assess the severity of loss of essential gene products to porcine cells, we investigated the rate at which gRNAs, and by association cells lacking genes targeted by these guides, were depleted in our GeCKO cell populations. To achieve this we clustered gRNA abundance profiles from PK15 cells (Figure 4a) and grouped them into three categories: (i) highly essential – genes that are depleted by day 5 (Figure 4b, cluster 1); (ii) mildly essential – genes that are depleted gradually over time (Figure 4b, cluster 2); and (iii) moderately essential – genes that are depleted between days 5 and 13 (Figure 4b, cluster 3). We found that most of our genes were classified as moderately essential (1,299 genes), with the majority shared in humans (64.3%, Figure 4c). These genes were spread across core cell function pathways involved in protein ‘Folding and degradation’, ‘DNA replication and repair’, ‘Transcription’, and ‘Translation’ (Figure 4d). Of the 527 highly essential genes, about a third (33.4%) were shared with humans. Interestingly the majority of essential lncRNAs (69.5%) also belonged to this category. These genes were mostly enriched in ‘DNA replication and repair’, ‘Transcription’, and ‘Translation’ (Figure 4d). Mildly essential genes had the fewest essential genes shared with humans (29.2%, Figure 4c), and the enriched pathways did not generally overlap with those of highly and moderately essential genes (Figure 4d).

**Figure 4:**
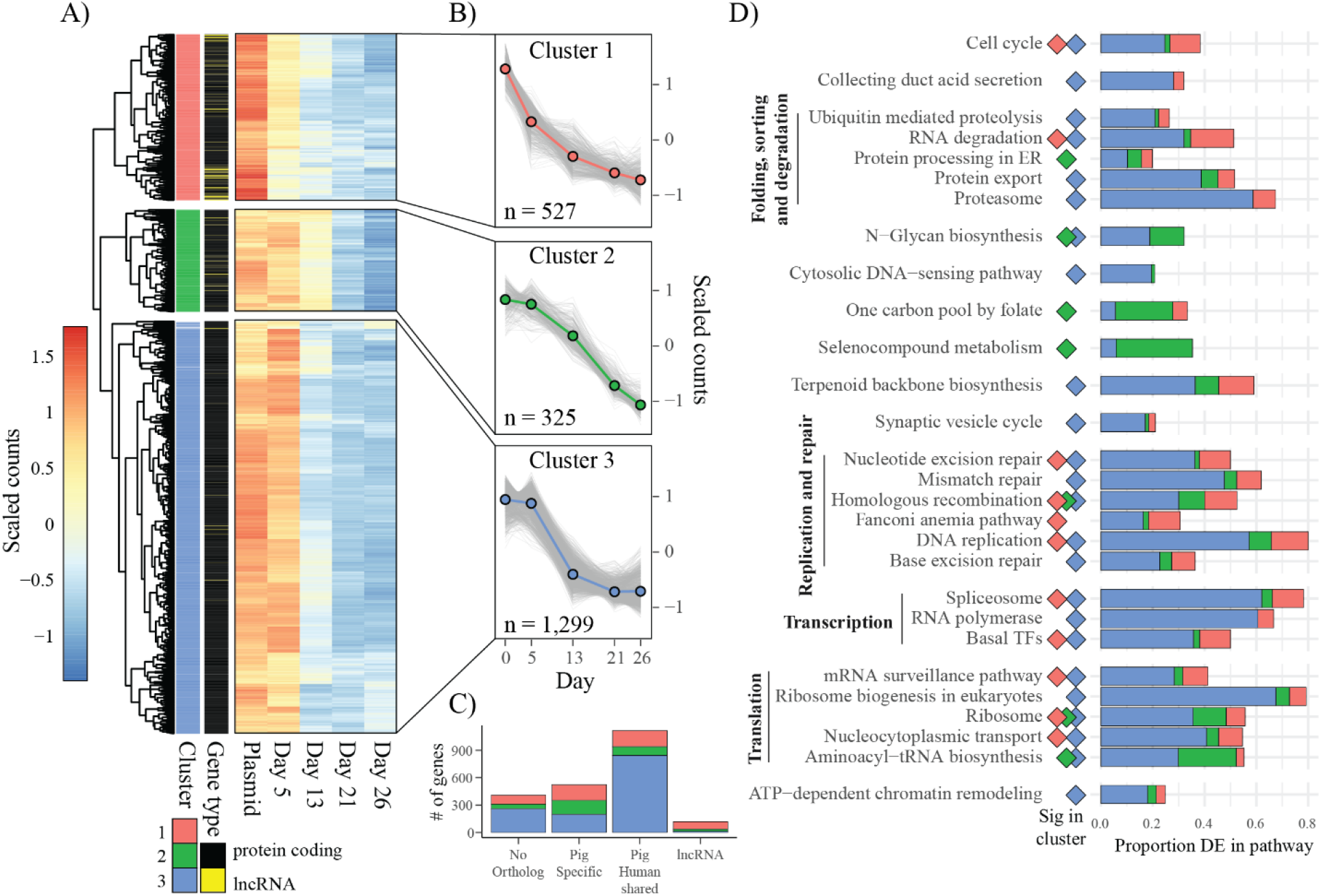
Rate of gene target depletion over time in PK15 cells. A) Heatmap of mean scaled gRNA counts for essential genes in PK15 cells. Genes are broken into three clusters based on trend of gRNA abundance over time. Genes with a correlation to the cluster mean < 0.8 were excluded from the cluster. Genes are annotated as either protein coding (black) or lncRNA (yellow). B) Cluster trends of gRNA abundance over time. Colored line corresponds to mean cluster trend while grey lines correspond to mean scaled gRNA abundance of each gene. C) KEGG enrichment of genes within each cluster.

## 5 Discussion

To our knowledge, this is the first study to systematically identify essential genes across the porcine genome. As expected, most of the essential genes we identified in our screen belonged to biological pathways core to cellular function, many of which are already known to be essential in human cells. Previous work identifying essential genes has shown that they can either belong to a core group that is essential in all cells or a conditional group that is specific to certain tissues, life-stages, and cell lines (Dvir et al., 2022). We find this to be the case in our study as well, with many essential genes in pigs being non-essential in humans, and some being cell line– or tissue-specific. In our study we found many more essential genes in PK15 cells than IPECJ2 cells. This discrepancy is likely due to bottlenecking of cell populations during passaging. For PK15 cells, we transferred around 10x more cells each passage than for IPECJ2 cells. This resulted in the screen in IPECJ2 cells to be statistically underpowered compared to the PK15 screen. As a result, it was difficult to differentiate between core and conditional essential genes in PK15 cells, as many of these were likely not statistically significant due to insufficient power in IPECJ2 screen. However, we did find 236 genes essential in IPECJ2 cells that are not essential in PK15 cells, suggesting that these genes are conditionally essential in IPECJ2 cells, as they were not identified in the highly powered PK15 screen.

Identification of embryonic lethal haplotypes in domestic pig populations is a high priority for animal breeders and several studies have identified lethal candidate genes (Derks et al., 2019; Derks et al., 2018; Derks et al., 2017; Derks et al., 2021). Among these, Derks et. al. (2019) identified four genes (*POLR1B, TADA2A, URB1*, and *PNKP*) as causal for homozygous embryonic lethality. In our screen we identify *POLR1B, URB1*, and *PNKP* as being essential for cell survival in both cell lines. Embryonic lethal genes in other studies included *BBS9* and *BMPER* (Derks et al., 2018); *KLHL40, POMGNT2, UROS*, and *ADAM12* (Derks et al., 2017); and *MYO7A* (Derks et al., 2021). However, none of these genes were essential in our screen, suggesting that they are essential during development but not for cell survival *in vitro*.

We identified 11 novel essential lncRNAs in our screen that were significantly depleted in both cell lines. Long non-coding RNAs are a large class of RNA with a wide range of functions, including transcriptional, post-transcriptional, epigenetic, and translational regulation (Mattick et al., 2023). In humans at least 173 essential lncRNAs have been identified (Zhang et al., 2022). Of the 11 essential lncRNAs that we found, six were annotated to regions enriched in rRNA genes on unplaced scaffolds, with three overlapping rRNA genes. In humans, ribosomal RNA biosynthesis is known to be highly regulated by lncRNAs through various mechanisms, such as the formation of heterochromatin using pRNA and the regulation of rRNA transcription through cryptic polII transcribed 5S-OT lncRNAs (Yan et al., 2019). It is possible that these lncRNAs are involved in regulation of rRNA synthesis through one of these mechanisms, or through a yet undescribed pathway. We also identified two essential lncRNAs on chromosome 10 which are transcribed in opposite directions with overlapping transcripts at their 3’ ends. Interestingly, these lncRNAs appear to have homology to centromere protein B which plays a key role in centromere formation (Sullivan & Glass, 1991).

The use of GeCKO screens in production species is a relatively new development, with the first porcine specific GeCKO library being used to identify host factors against Japanese encephalitis virus in 2020 (Zhao et al., 2020). Since then, porcine specific screens have been employed to identify host genes influencing infection of pig cells with influenza (Zhou et al., 2021), porcine reproductive and respiratory syndrome (PRRSV) (Jiang et al., 2022), and African swine fever (Shen et al., 2024). The largest of these libraries was for African swine fever, containing 186,510 gRNAs targeting all porcine coding genes, lncRNAs, and miRNAs with a redundancy of 6-10 guides. Our library similarly targets all porcine coding genes, lncRNAs, and miRNAs, but with a lower redundancy of 3-5 guides per element. Despite this, we were able to identify essential genes with fewer cells, achieving a guide representation of over 400 cells per guide. This approach makes GeKCO experiments more manageable while still maintaining sufficient guide representation for statistical significance.

## 6 Conclusion

This study provides a significant advancement in our understanding of essential genes within the porcine genome. By using a novel pig-specific GeCKO library, we identified 2,281 and 928 essential genes in PK15 and IPEC-J2 cells, respectively. Our findings underscore the importance of core cellular processes such as cell cycle regulation, DNA replication, transcription, and translation, highlighting the conservation of essential genetic elements across species. Comparative analysis with human essential genes from the DepMap project revealed a substantial overlap, with over half of the identified porcine essential genes being conserved in humans. Additionally, the study uncovered porcine-specific essential genes and lncRNAs, providing new insights into unique aspects of porcine molecular physiology.

Overall, this work establishes a strong foundation for future studies aimed at leveraging porcine models to enhance our understanding of human diseases and develop therapeutic interventions. By elucidating the landscape of essential genes in pigs, this research not only deepens our understanding of porcine genetics but also contributes to the broader goal of improving human health through more accurate and translatable animal models.

