## Supplementary figures and images for "Genome-wide CRISPR knockout screen reveals the landscape of essential genes across the porcine genome"

### Figure S1

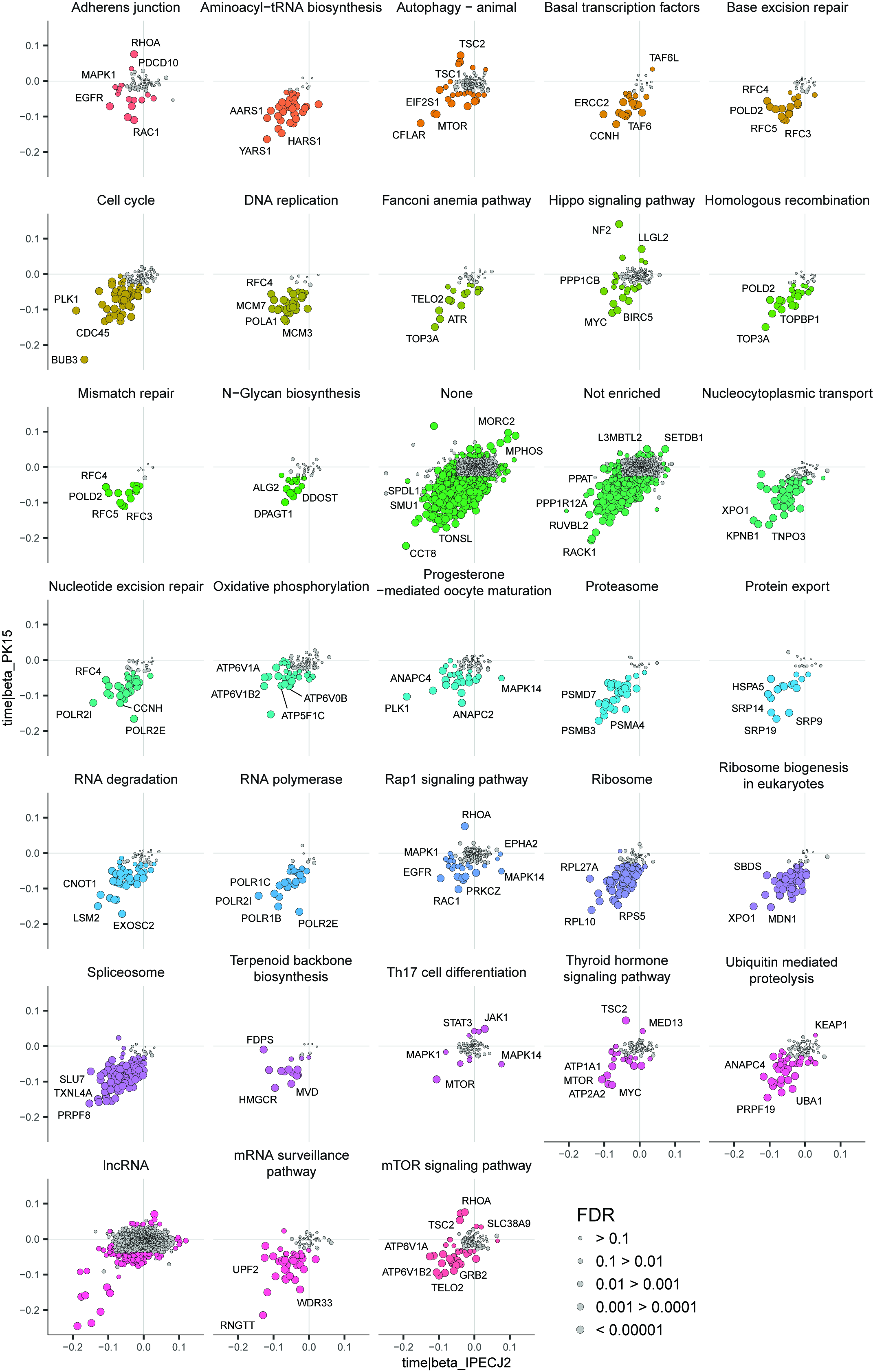
